# Insights into the nutritional properties and microbiome diversity in sweet and sour yogurt manufactured in Bangladesh

**DOI:** 10.1101/2021.08.15.456382

**Authors:** S. M. Rafiqul Islam, Afsana Yeasmin Tanzina, Md Javed Foysal, M. Nazmul Hoque, AMAM Zonaed Siddiki, Alfred Tay, S. M. Jakir Hossain, Muhammad Abu Bakar, Mohammad Mostafa, Meheadi Hasan Rumi, Adnan Mannan

**Affiliations:** Department of Genetic Engineering and Biotechnology, Faculty of Biological Sciences, University of Chittagong, Chattogram-4331, Bangladesh; Department of Genetic Engineering and Biotechnology, Shahjalal University of Science and Technology, Sylhet-3114, Bangladesh; Department of Gynecology, Obstetrics and Reproductive Health, Bangabandhu Sheikh Mujibur Rahman Agricultural University, Gazipur-1706, Bangladesh; Department of Pathology and Parasitology, Chattogram Veterinary and Animal Sciences University, Chattogram-4225, Bangladesh; Helicobacter Research Laboratory, Marshall Centre for Infectious Disease Research and Training, School of Biomedical Sciences, University of Western Australia, Perth, WA, Australia; Laboratory of Forest Chemistry, Bangladesh Forest Research Institute, Chattogram-4211, Bangladesh; BCSIR Laboratories, Chattogram-4220, Bangladesh

**Keywords:** Yogurt, Brands, Tastes, Metagenomics, Nutritional contents, Microbiome diversity

## Abstract

Yogurt quality mainly depends on nutritional properties, microbial diversity and purity of starter culture. This study aimed to assess the nutritional composition and microbiome diversity in yogurt. Microbial diversity was analyzed by 16S and 18S rRNA based high-throughput sequencing. Significantly (P<0.05) higher pH, fat, moisture, total solid and solid-non-fat contents (%) were observed in sweet yogurt whereas sour varieties had significantly higher ash and minerals. Metagenomic investigation showed that 44.86% and 55.14% reads were assigned to bacterial and fungal taxa, respectively, with significantly higher taxonomic richness in sour yogurt. A significant difference in bacterial (P_permanova_=0.001) and fungal (P_permanova_=0.013) diversity between sweet and sour yogurt was recorded. We detected 76 bacterial and 70 fungal genera across these samples which were mostly represented by Firmicutes (>92%) and Ascomycota (98%) phyla, respectively. Among the detected genera, 36.84% bacterial and 22.86% fungal genera were found in both yogurt types. Our results suggest that *Streptococcus* (50.82%), *Lactobacillus* (39.92%), *Enterobacter* (4.85%), *Lactococcus* (2.84%) and *Aeromonas* (0.65%) are the most abundant bacterial genera, while *Kluyveromyces* (65.75%), *Trichosporon* (8.21%), *Clavispora* (7.19%), *Candida* (6.71%), *Iodophanus* (2.22%), *Apiotrichum* (1.94%), and *Issatchenkia* (1.35%) are the most abundant fungal genera in yogurt metagenomes. This is the first study on nutritional properties and microbiome diversity of Bangladeshi yogurt that would be a benchmark for safe production of quality yogurt by commercial manufacturers.

## Introduction

Dairy products are a potential source of nutrients for humans that provide necessary energy, calcium, protein, and other micro- and macronutrients^1^. Fermented dairy products are associated with numerous health benefits while reducing the risks of different diseases or disorders including obesity prevention, suppression of metabolic diseases and disorders, reduction of possible risks of cardiovascular and immune-related diseases through providing the consumer with both readily metabolizable nutrients and beneficial microorganisms^2,3^. Worldwide more than 400 diversified fermented products are manufactured from milk and yogurt is one of the major universal fermented dairy food which is locally well known as “Dahi” in Indian subcontinent^4^. Milk is a major nutritional resource^5^ and presents a complex ecosystem of interconnected microbial communities which modulates both health and diseases^6,7^. Yogurt is a nutrient-rich food that provides the consumers with necessary source of protein, calcium, magnesium, vitamin B12 and some key fatty acids^8^. Moreover, yogurt is used as a major source of probiotics, which are beneficial bacteria thought to improve human health^4^.

Yogurt, prepared from cows and buffaloes milk, is a rich source of diverse microbial communities, which indeed could vary within the yogurt of different tastes (sweet and sour) and different manufacturers. Understanding the actual composition of the yogurt microbial community (bacteria and fungi), and its relevance with the quality and safety of yogurt, is of paramount importance. The microbiome diversity and composition of the yogurt can thereby regulates the development of the organoleptic properties of yogurt, nutrient composition, shelf-life and associated safety^9^. In Bangladesh, various fermented milk products including yogurt, lassi, mattha, borhani etc. are produced as a pattern of ethnicity. Among these, yogurt is the most popular due to its sensory attributes along with nutritional value and positive impacts in promoting digestion and gastrointestinal disorders^10^. However, these beneficial effects of yogurt depend on both qualitative and quantitative composition of the constituent microflora in the yogurt which need to be ascertained. Traditionally in Bangladesh, yogurt is manufactured by fermentation of cow and buffalo milk or a mixture of them at an optimal pH 4.0–4.6 primarily through the action of *Streptococcus salivarius* subsp. *thermophilus*, and *Lactobacillus delbrueckii* subsp. *bulgaricus*^11^.

According to FAO (Food and Agriculture Organization), regular yogurt contains 5–6% protein, 8.25% solid-not-fat (SNF), and 3.5% fat, though the fat content varies from 0–3.5% based on the type of yogurt^12,13^. Yogurt is a good source of protein, vitamins (including vitamin A, vitamin B2, vitamin B5 and vitamin B12), minerals (including sodium, potassium, calcium, magnesium, iron, zinc and copper) and some key fatty acids^13,14^. Though the conventional starter culture of yogurt includes only *S. thermophilus* and *L. bulgaricus*, its characteristic pH favors the growth of other microorganisms (e.g., *Lactococcus lactis* subsp. *diacetylactis*, *Lactococcus. cremoris* and *Lactobacillus rhamnosus*) which contribute to yogurt viscosity, appealing aroma and taste, and thought to have probiotic effects^4,15^. The major drawback of these conventional culture-based screening methods is its inability to reveal microbes that are insensitive to culture but responsible for a greater impact on fermented foods.

Recent advances in culture-independent techniques such as high-throughput sequencing (HTS) and next generation sequencing (NGS) technology along with advanced bioinformatics and computational tools during the last decade have revolutionized the research on food microbial ecology, leading to consider microbial populations as consortia^7,16,17^. By employing this technology, unculturable microbial flora including both beneficial and pathogenic bacteria, as well as fungi have been successfully identified in various naturally fermented milk (NFM) products ^17,18–19^. One of the recent studies conducted using both culture-dependent and culture-independent methods reported *Lactobacillus* and *Streptococcus* as dominant bacteria and *Kodamaea*, *Clavispora*, *Candida*, and *Tricosporon* as dominant fungal genera in four traditional Bangladeshi fermented milk products (dahi, chanar-misti, paneer, and borhani)^20^. Another study conducted by Rashid et al.^21^ identified bacterial species including *L. bulgaricus*, *L lactis*, *L. fermentum*, *S. thermophilus*, *S. bovis*, *E. faecium*, *L. mesenteroides*, *L. dextranicum*, *Lc. lactis*, *Lc. raffinolactis*, *P. pentosaceus* in yogurt from different regions. However, these studies did not indicate any characteristic microbial features of a particular brand or taste types.

Like milk and other dairy products, yogurt has been reported to be a unique source of trace minerals Na, K, Ca, Mg, Fe, Zn and Cu^22,23^. The content of trace elements in dairy milk and products has begun to be more widely studied, particularly in industrialized and polluted regions, since it is considered a good bioindicator of pollution of the agricultural environment^24–26^. However, to our knowledge, no comprehensive research has been carried out in Bangladesh as of now for investigating the presence of nutritional elements and trace minerals in yogurt. Besides, detailed knowledge on biochemical potential and variations in microbiome composition along with diversity among different brands and taste types is yet to be analyzed. In this study, we aimed to assess the quality of sweet and sour yogurt samples of seven reputed brands produced from cow and buffalo milk and detect yogurt microbiome diversity. We conducted this study in two stages; in stage one, the biochemical parameters and trace minerals of the yogurt samples were determined to ensure its nutritional status and safety and in stage two, we scrutinized the microbial (bacteria and fungi) community through targeted sequencing of the 16S and 18S rRNA genes via the Illumina MiSeq sequencing platform for uncovering the microbial consortia and their activities in yogurt.

## Results

### Nutritional composition of yogurt

#### Analysis of physical and biochemical parameters

The quality and safety of yogurt depends on its nutritional composition and microbiome diversity as well as quality of the starter culture. To assess yogurt quality, the nutritional properties of yogurt in terms of biochemical parameters and trace minerals were examined in this study. Biochemical parameters of yogurt samples including pH, fat, moisture, total solid (TS), solid-not-fat (SNF) and ash contents are summarized in Table 1. During the commercial yogurt manufacturing process, pH needs to be maintained so that the taste doesn’t become rancid and the flavor is proper. A high pH indicates improper fermentation and allows many undesirable microorganisms to grow in the medium. We found significant differences (*P* < 0.05) in pH values of all yogurt varieties studied in this report. The mean pH values of the yogurt samples remained slightly acidic and, on an average, ranged from 5.28–6.33. The sweet yogurt Brand 6 (SwV5) had the highest average pH value of 6.33±0.13 whereas the sour yogurt Brand 7 (SoV5) showed the lowest pH (5.28±0.25) (Table 1). Interestingly, no significant difference was found for fat content of the analyzed yogurt samples. The average fat content of the samples varied from 0.25 to 2.52% (w/w). The sweet yogurt of Brand 1 (SwV1) had the highest fat content (2.52±3.68) while the lowest fat content (0.25±0.05) was recorded in sour yogurt Brand 3 (SoV2) (Table 1).

**Table 1.** Biochemical parameters of Bangladeshi yogurt of different brands and tastes.

| Brands | Parameters (% w/w; except pH) |  |  |  |  |  |
| --- | --- | --- | --- | --- | --- | --- |
|  | pH | Fat<sup>A</sup> | Moisture | TS | SNF | Ash<sup>A</sup> |
| Brand 1<br>(SwV1) | 5.8<sup>ab</sup>±0.16 | 2.52±3.68 | 82.1<sup>a</sup>±1.16 | 17.90<sup>a</sup>±1.16 | 15.37<sup>a</sup>±3.36 | 0.86±0.23 |
| Brand 2<br>(SwV2) | 6.2<sup>a</sup>±0.04 | 0.93±0.30 | 78.88<sup>b</sup>±0.38 | 21.11<sup>b</sup>±0.38 | 20.18<sup>b</sup>±0.58 | 0.93±0.30 |
| Brand 2<br>(SoV1) | 5.84<sup>ab</sup>±0.11 | 1.41±0.66 | 82.44<sup>a</sup>±0.69 | 17.56<sup>a</sup>±0.69 | 16.16<sup>a</sup>±0.80 | 1.06±0.30 |
| Brand 3<br>(SwV3) | 6.32<sup>a</sup>±0.17 | 0.9±0.62 | 80.53<sup>ab</sup>±0.84 | 19.47<sup>ab</sup>±0.84 | 18.57<sup>ab</sup>±1.13 | 1.13±0.31 |
| Brand 3<br>(SoV2) | 5.71<sup>ab</sup>±0.09 | 0.25±0.05 | 87.59±0.53 | 12.41±0.53 | 12.16<sup>a</sup>±0.51 | 1.06±0.31 |
| Brand 4<br>(SwV4) | 6.28<sup>a</sup>±0.01 | 0.8±0.2 | 60.72±0.80 | 39.29±0.80 | 38.48±0.74 | 0.80±0.20 |
| Brand 4<br>(SoV3) | 5.98<sup>a</sup>±0.03 | 2.15±1.91 | 79.17<sup>b</sup>±0.49 | 20.82<sup>b</sup>±0.49 | 18.67<sup>ab</sup>±1.46 | 1.26±0.31 |
| Brand 5<br>(SoV4B) | 6.28<sup>a</sup>±0.01 | 1.6±0.2 | 84.74±0.64 | 15.26±0.64 | 13.66<sup>a</sup>±0.53 | 1.13±0.42 |
| Brand 6<br>(SwV5) | 6.33<sup>a</sup>±0.13 | 2.32±1.80 | 75.83±0.73 | 24.17±0.73 | 21.85<sup>bc</sup>±1.07 | 0.80±0.20 |
| Brand 7<br>(SoV5) | 5.28±0.25 | 2.19±0.19 | 72.06±0.82 | 27.94±0.82 | 25.75<sup>c</sup>±1.01 | 1.0±0.20 |
| Level of<br>sig. | *** | NS | *** | *** | *** | NS |

*TS = Total solid, SNF = Solid-not-fat, SwV = Sweet variety, SoV = Sour variety, Mean values with different* *superscripts (a, b, c, ab, bc) in the same column differed significantly while those left with no superscripts differed* *with all, A = No mean value showed any significant difference among each other hence were left single i.e. without* *any superscripts, \*\*\* = Significant at 5% level (P < 0.05), NS = Non-significant, The values of all parameters* *were recorded as mean ± SD.*

Moisture content (MC) is one of the most common properties of food products due to a number of reasons like food quality, microbial stability along with legal and labeling requirements. The MC of the yogurt samples were in the range of 60.72±0.80 to 87.59±0.53% and varied significantly (*P* < 0.05) across the sample groups (Table 1). The average MC was much higher for all sweet Brands compared to sour Brands except for SwV4. The sour varieties of Brand 3 (SoV2) had the highest MC (87.59±0.53%) and lowest MC (60.72±0.80%) was recorded in sweet varieties of Brand 4 (SwV4). Likewise, the TS and SNF values differed significantly (*P* < 0.05) across the sample groups. We found the highest amount of TS (39.29±0.80%) and SNF (38.48±0.74%) in sweet Brand 4 (SwV4) yogurt which remained lowest (TS; 12.41±0.53% and SNF; 12.16±0.51%) in sour Brand 3 (SoV2) yogurt (Table 1). However, after removing the organic residues present in samples, Brand 4 (SwV4) and Brand 6 (SwV5) yogurt samples yielded the lowest ash content (0.80±0.20%), while the highest amount of ash content (1.26±0.31%) was found in sour varieties of Brand 4 (SoV3) yogurt. No significant difference in ash content was found among the sample varieties.

#### Analysis of mineral contents

Yogurt is an important source of essential minerals. Yet some toxic elements (Cr, Ni and Cd) may be accidently added during handling, processing, and remixing of milk to prepare yogurt. The contents of minerals including Na, K, Ca, Mg, Fe, Zn, and Cu in yogurt samples of different brands and tastes are given in Table 2. The mean value for Na content ranged from 593.50±65.55 to 1052.65±332.42 mg/kg indicating its highest and lowest content in sour yogurt Brand 4 (SoV3) and Brand 3 (SoV2), respectively. The Brand 4 (SoV3) of sour yogurt had the highest content of K (2744.54±669.79 mg/kg), Ca (2442.27±92.21 mg/kg), and Mg (272.42±27 mg/kg). The lowest concentration of K (719.36±135.50 mg/kg) was found in another sour Brand 5 (SoV4B), while the sweet yogurt Brand 4 (SwV4) had the least mean content of Ca (1115.40±354.57 mg/kg) and Mg (107.72±20.05 mg/kg). Among these trace elements, the value of K varied within a broad range of 719.36±135.5 – 2744.52±669.79 mg/kg.

**Table 2.**
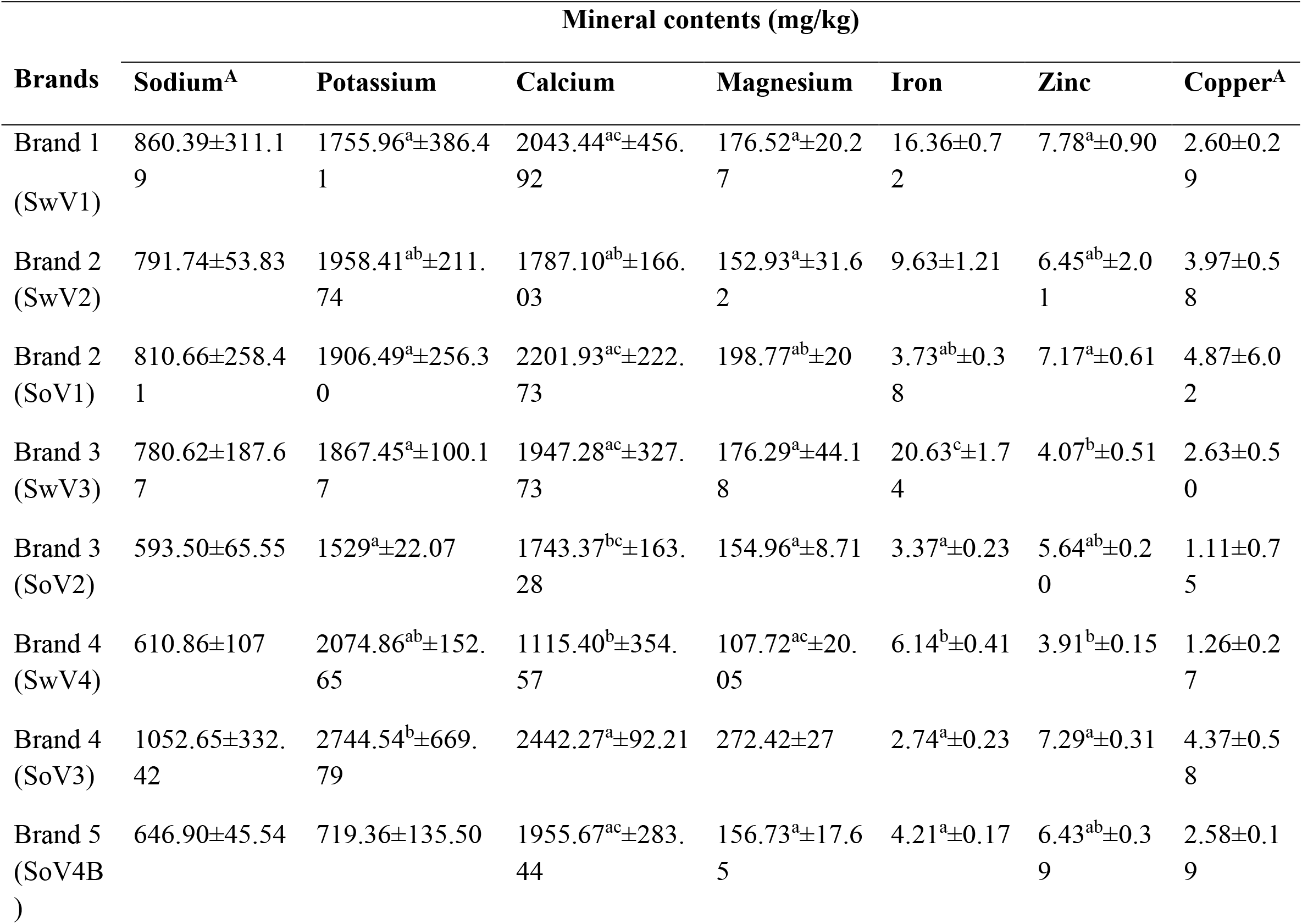
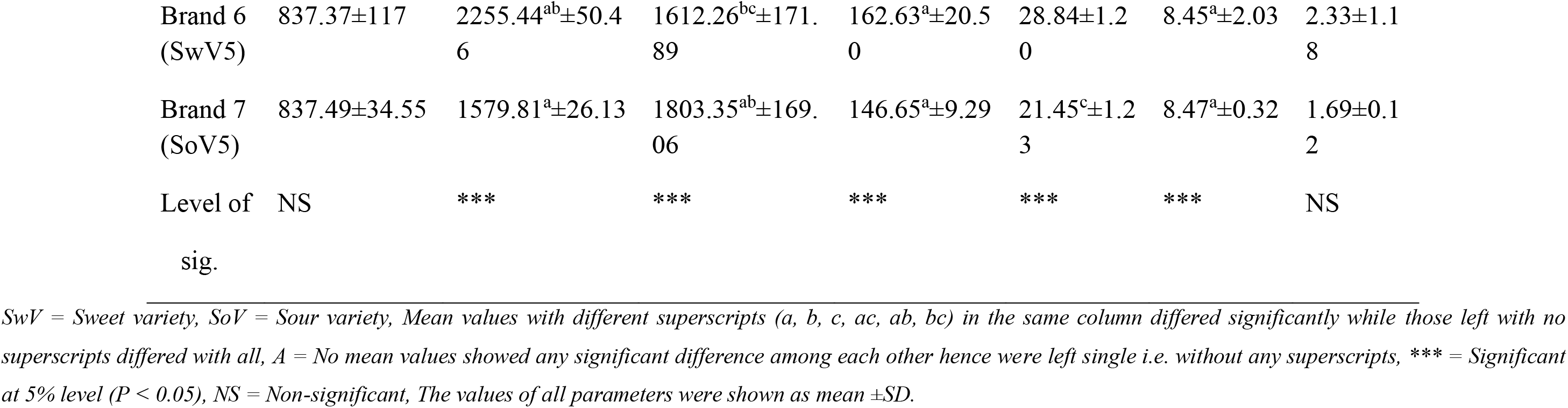
The mineral contents in Bangladeshi yogurt of different brands and tastes (sweet and sour).

The concentration of Fe in yogurt varied from 2.74±0.23 to 8.84±1.20 mg/kg. The content of Fe in different brands of yogurt was statistically significant (p < 0.05) since the sweet yogurt always had higher mean values of Fe content except for one sour sample (SoV5) of Brand 7 in which the Fe content was detected as 21.45±1.23 mg/kg (Table 2). The concentration of Zn ranged from 3.91±0.15– 8.47±0.32 mg/kg. The highest accumulation of Zn (8.47±0.3 mg/kg) was noted in sour yogurt of Brand 7 (SoV5) followed by in sample SwV5 of Brand 6 (8.45±2.03 mg/kg), while the least concentration (3.91±0.15 mg/kg) was detected in Brand 4 of sweet yogurt (SwV4). The amount of Cu varied within the range of 1.11±0.75 to 4.87±6.02 mg/kg. The maximum value was detected in SoV1 of sour yogurt Brand 2 with the lowest content of Cu (1.11±0.75 mg/kg) in SoV2 of Brand 3. The extreme SD value with the maximum level of Cu indicates the concentration of Cu differed significantly (*P* < 0.05) among the brands of yogurt (Table 2). These results highlighted that mineral contents vary among different commercial brands (brand1–brand7) and taste types (sweet and sour) of yogurt. Although minerals represent a small portion of yogurt, they are fundamental for human health, yogurt quality and its characteristic tastes.

### Microbiome composition and diversity

Yogurt microbiomes of 30 samples (sweet = 15 and sour = 15) belonging to seven different commercial brands (Brand 1– Brand 7) were analyzed through high-throughput amplicon sequencing. The sweet yogurt samples included only cows samples (n = 15), and the sour yogurt included both cows (n =12) and buffaloes (n = 3) samples. During this study, the targeted sequencing approach generated a total of 3.10 million high quality reads (with an average of 0.104 million reads per sample), of which 1.4 and 1.7 million reads were assigned into 306 bacterial and 3144 fungal OTUs (Data S1). Among the reads, 44.86% and 55.14% reads were assigned to bacterial and fungal taxa, respectively. There is a clear difference between phylogenetic profiles and microbiota quantitation obtained using 16S rRNA (V3-V4) and ITS primers. An average good’s coverage index of 0.995 for bacteria and 0.996 for fungi indicated that sequencing depth was sufficient enough to capture most of the microbial community at different taxa levels (phylum, order, genus etc.) (Data S1).

The alpha-beta diversity of microbiomes was analyzed to observe the differences in microbial composition and diversity in yogurt. Fig. 1 shows the alpha-beta diversity of bacteria and fungi communities in yogurt samples of different brands and tastes types (sweet and sour). In this study, we estimated both within sample (alpha) and between sample (beta) diversities in yogurt samples of seven different brands (Brand 1–7) and two different tastes (sweet and sour) using different diversity indices (Fig. 1). We found significant differences in α-diversity (observed richness, Shannon and Simpson estimated) in bacterial components of the microbiomes across the samples of both sweet and sour yogurts, and different brands (Fig. 1A, B). This diversity difference was evidenced significantly by higher taxonomic resolution in sour yogurt especially in Brand 5 (Fig. 1A). Though, the observed species and Shannon estimated diversity of the fungal fraction of the yogurt microbiomes varied among different brands, however remained less diverse between the taste types (sour and sweet) of yogurt (Fig. 1C, D). Moreover, the α-diversity of the fungal component of the yogurt microbiomes was found more diverse in sour yogurt compared to sweet yogurt samples (Fig. 1D).

**Fig. 1.**
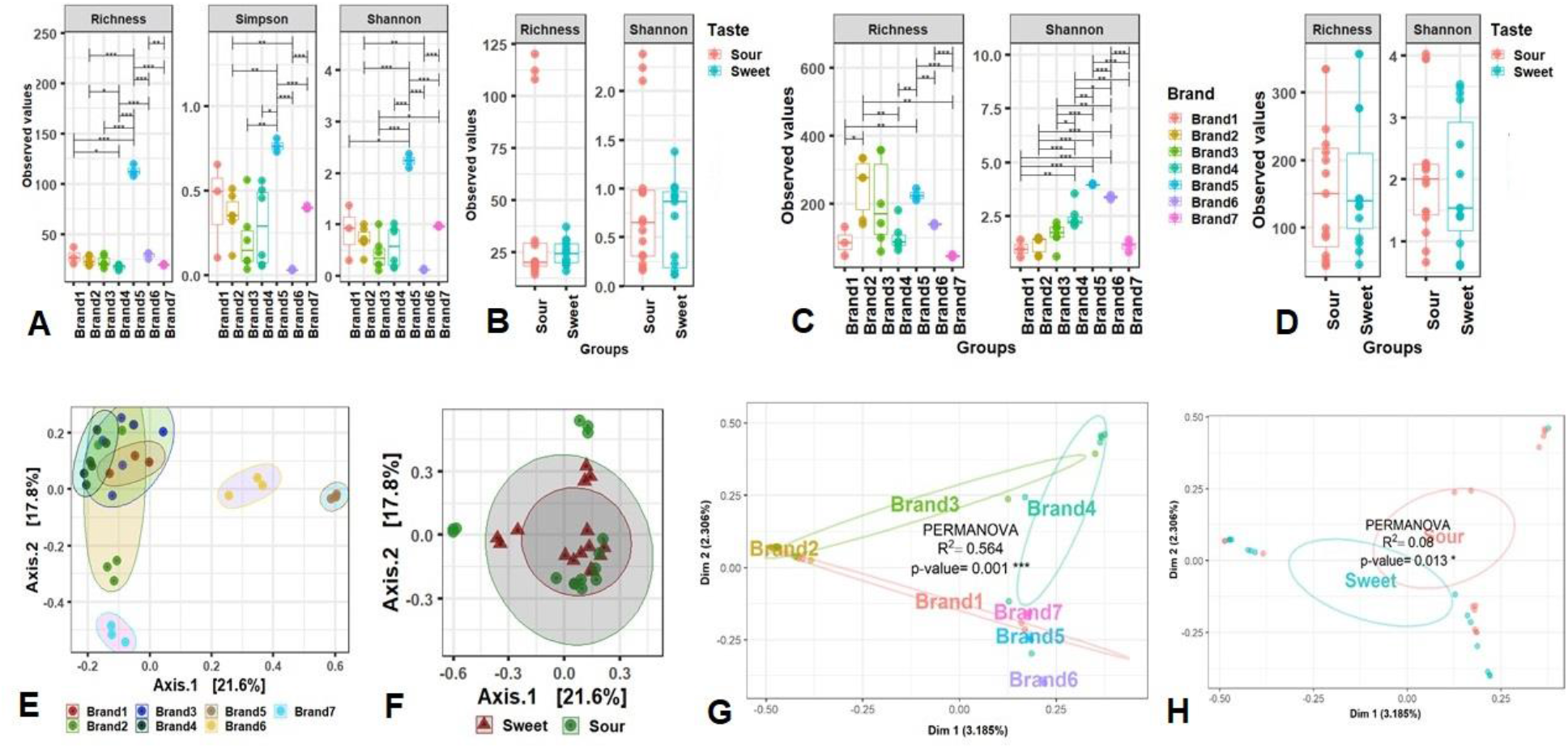
Differences in microbiome diversity and community structure in seven different brands (Brand 1–7) and two different tastes (sweet and sour) of yogurt samples. Alpha diversity was measured through the observed species richness, Simpson and Shannon estimated diversity indices. **(A, B)** α-diversity of bacterial communities and **(C, D)** α-diversity of fungal fraction of yogurt microbiomes; **(E, F)** Principal coordinate analysis (PCoA) to measure the β-diversity in bacterial component of the yogurt microbiomes segregated yogurt samples according to **(E)** seven different brands and **(F)** two different tastes (sweet and sour); **(G, H)** Beta ordination (NMDS) plots representing the fungal component of the microbiomes separated yogurt samples according to **(G)** different brands and **(H)** two different tastes (sweet and sour) categories. Statistical analysis using Kruskal-Wallis tests showed significant microbial diversity variation (P < 0.01 = *, P < 0.001 = ***).

The principal coordinate analysis (PCoA) plots representing β-diversity showed significant differences (*P* = 0.001, Kruskal-Wallis test) in bacteriome composition among different brands (Fig. 1E). We also found significant differences (*P* = 0.001, Kruskal-Wallis test) in bacterial diversity between sweet and sour yogurt sample groups (Fig. 1F), representing the prevalence of different sets of bacteria in each sample category. Similarly, NMDS estimated beta-ordination plots depicting the diversity of the fungal fraction of the yogurt microbiomes also showed significant variations in different brands (*P*_permanova_ = 0.001) and tastes (*P*_permanova_ = 0.013) (Fig.1G, H). For instance, Brand 3 maintained the highest beta-dispersion in relation to Brand 6, Brand 5, Brand 2, and Brand 7, respectively (Table S1). Least significant clustering differences were detected between the Brand 1 and 6, and Brand 1 and 2 (Fig. 1G).

### Multivariate analysis and downstream bioinformatics

During data analyses, the detected OTUs were assigned into 11 phyla and 76 genera for bacteria, and 5 phyla and 70 genera for fungi. We detected 11 bacterial phyla including 10 in sweet yogurt of cow milk, 10 and 9 in sour yogurt samples of cow and buffalo milk, respectively (Table S2). Among the identified bacterial phyla, *Firmicutes* was the most abundant phylum with a relative abundance of 92.89% followed by *Proteobacteria* (7.03%) (Fig. S2A). By comparing the relative abundances of these predominant phyla, we found that *Firmicutes* was the most abundant phylum in both sweet (99.84%) and sour (86.36%) yogurt samples of cow milk. Conversely, *Proteobacteria* was the most abundant phylum in sour yogurt (61.46%) of buffalo milk followed by *Firmicutes* (38.12%) (Fig. S2A, Data S1). In addition, we identified 30 orders of bacteria across these sample groups, of which 24, 25 and 21 orders were detected in sweet yogurt of cow, sour yogurt of cow and sour yogurt of buffalo, respectively (Table S2). Among the bacterial orders, *Lactobacillales* was found as the predominating order in sweet yogurt of cow (99.48%) and sour yogurt of cow (86.20%), while the sour yogurt of buffalo milk had higher abundance of *Enterobacteriales* (52.82%), *Lactobacillales* (37.48%) and *Aeromonadales* (5.52%) (Data S1).

The taxonomic classification and comparison of microbiomes at genus-level in yogurt samples of different tastes and milk sources are illustrated in Fig 2. In this study, the yogurt samples collectively harbored 76 bacterial genera (Fig. 2A), and of them, *Streptococcus* (50.82%), *Lactobacillus* (39.92%), *Enterobacter* (4.85%), *Lactococcus* (2.84%) and *Aeromonas* (0.65%) were the top abundant genera (Data S1). In addition, we demonstrated notable differences in the diversity and composition of the identified bacterial genera both in sweet and sour yogurt samples of different brands. Among the detected bacterial genera, 36.84% genera were found to be shared in sweet yogurt of cow, sour yogurt of cow and sour yogurt of buffalo metagenomes (Table S2). The sweet and sour yogurt of cow had a sole association of 7 (9.2%) and 4 (5.26%) genera, respectively (Fig. 2A, Data S1). Likewise, 70 fungal genera were detected in the study samples, of which 22.86% genera were shared in the metagenomes of sweet yogurt cow, sour yogurt cow and sour yogurt buffalo (Fig. 2B; Table S2). Of the detected fungal genera in both sweet and sour yogurt samples, *Kluyveromyces* (65.75%), *Trichosporon* (8.21%), *Clavispora* (7.19%), *Candida* (6.71%), *Iodophanus* (2.22%), *Apiotrichum* (1.94%), and *Issatchenkia* (1.35%) were the most abundant genera (Data S1). The sweet and sour yogurt samples of cow had a sole association of 16 (22.86%) and 18 (25.71%) fungal genera. However, the sour yogurt samples of buffalo had none of the bacterial and fungal genus found to be shared in the study metagenomes (Fig. 2B, Data S1).

**Fig. 2.**
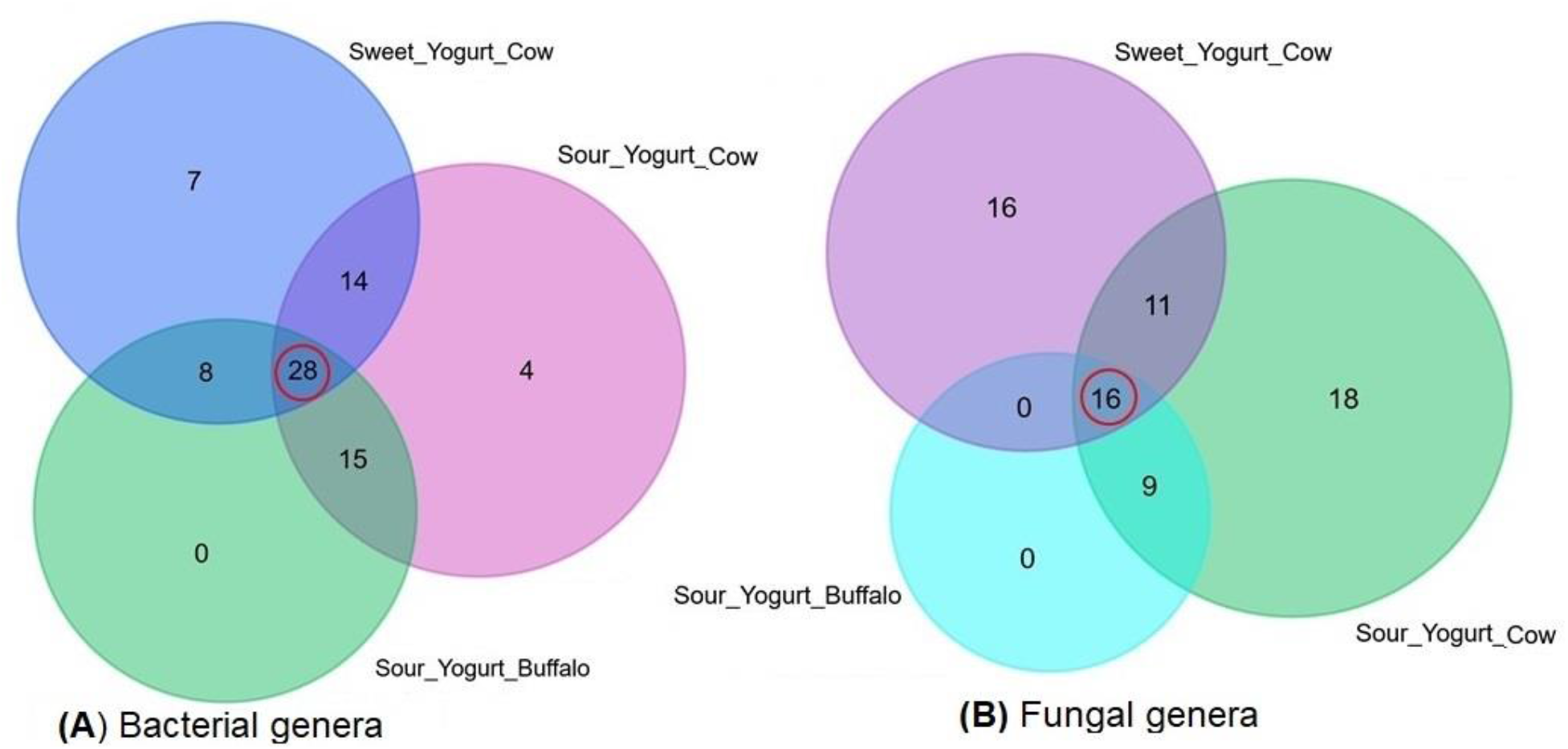
Taxonomic composition of microbiomes in two different tastes (sweet and sour), and milk sources (cow and buffalo) of yogurt samples. **(A)** Venn diagram showing unique and shared bacterial genera in sweet (cow) and sour (cow and buffalo) yogurt. Out of 76 detected genera, only 28 (36.84%) genera (highlighted in red circle) were found to be shared in sweet yogurt cow, sour yogurt cow and sour yogurt buffalo metagenomes. **(B)** Venn diagram comparison of unique and shared fungal genera identified in the study metagenomes. Of the detected fungal genera (n = 70), only 16 (22.86%) genera (highlighted in red circle) were found to be shared in the metagenomes of sweet yogurt cow, sour yogurt cow and sour yogurt buffalo samples. More information on the taxonomic composition and relative abundances is available in Data S1.

We also analyzed the taxonomic classification of bacteria and fungi at genus level in yogurt of various brands, tastes and milk source varieties (Fig. 3). Among the bacterial genera, *Streptococcus* (71.49%) and *Lactobacillus* (27.93%) had higher relative abundances in cow originated sweet yogurt samples (Fig. 3A; Data S1). On the other hand, sour yogurt samples of cow had higher relative abundances of *Lactobacillus* (50.34%), *Streptococcus* (30.32%), *Enterobacter* (9.28%), *Lactococcus* (5.41%) and *Aeromonas* (1.22%). Simultaneously, *Streptococcus* (42.43%), *Macrococcus* (24.76%), *Enterobacter* (11.88%), *Empedobacter* (9.76%), *Aeromonas* (5.54%) and *Enterococcus* (2.91%) were detected as the top abundant genera in sour yogurt of buffalo. The rest of the genera had relatively lower abundances (<1.0%) in these metagenomes. The heatmap representation (Fig. 4) of the yogurt microbiomes shows distinct separation among the detected bacterial genera into two major clusters according to their host origin i. e. cow and buffalo rather than tastes of the yogurt. The heatmap of relative abundance of bacteria and fungi at genus level across the tastes and source varieties of yogurt is also presented in Fig. 4. The cow originated bacterial genera of the yogurt microbiomes were further sub-clustered according to tastes (sweet and sour) (Fig. 4A, Data S1). Differential abundance of bacteria and fungi in yogurt samples of different brands and tastes is listed in Table 3. Differential abundance analysis also showed that *Aeromonas*, *Enterobacter*, *Lactococcus*, and *Acinetobacter* had significantly (Bonferroni corrected *P*_BC_ = 0.033–0.044, Bonferroni test) higher number of reads assigned for these genera in Brand 5 compared to other Brands (Table 3A). In addition, significantly (*P*_BC_ = 0.043, 0.024) higher number of reads assigned for *Streptococcus* and *Lactobacillus* were found in Brand 6 and 7, respectively. Significantly (*P*_BC_ = 0.028) greater number of *Streptococcus* associated reads (n = 4494.5) were observed in Brand 1 and this might be linked to the sweetness of yogurt brands (Table 3B).

**Fig. 3.**
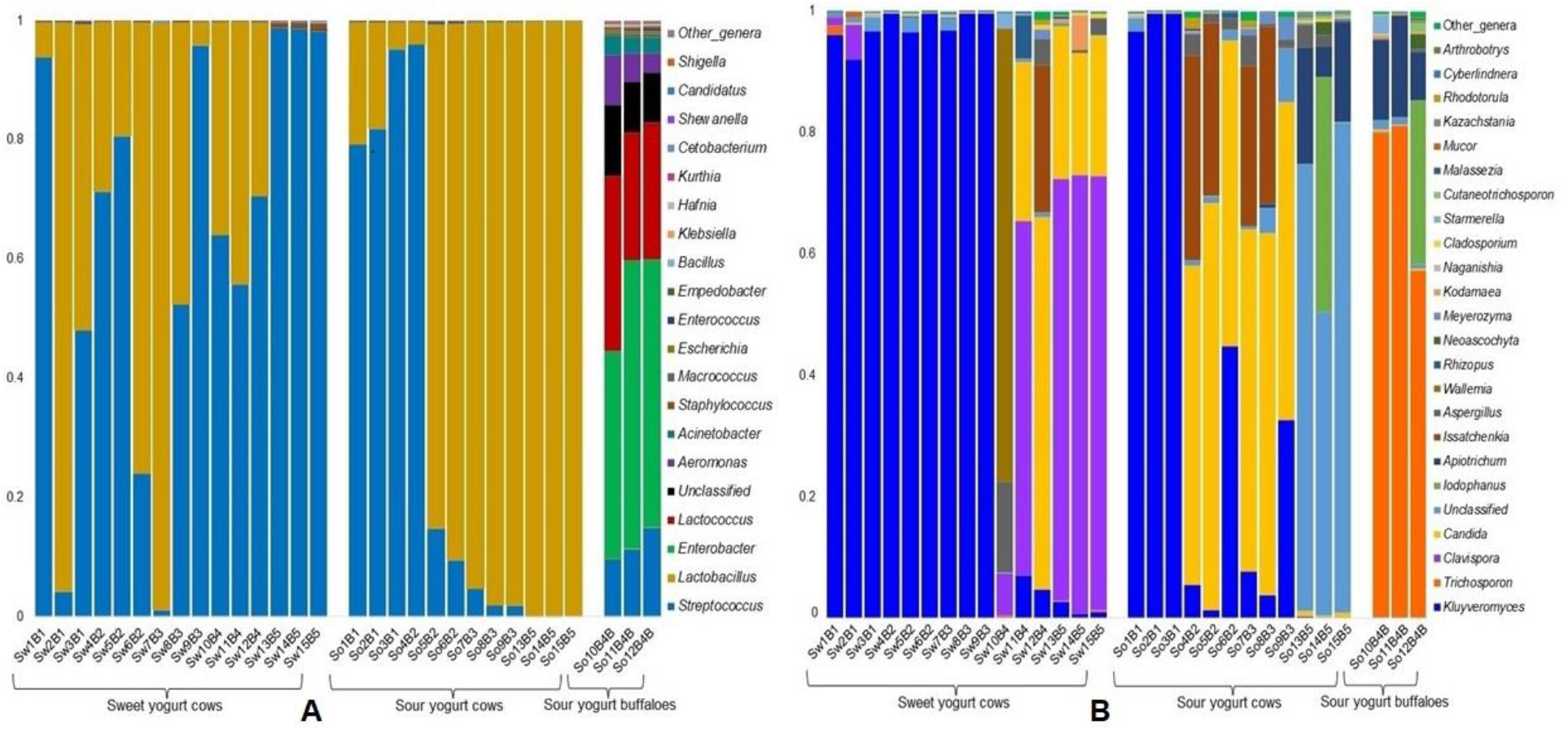
The genus level taxonomic profile of bacteria and fungi in sweet yogurt of cow, sour yogurt of both cow and buffalo samples. **(A)** The relative abundance of 20 most abundant bacterial genera in 30 samples is sorted from bottom to top by their deceasing proportion with the remaining genera keeping as ‘other genera’. **(B)** The relative abundance of 25 most abundant fungal genera in 30 samples is sorted from bottom to top by their deceasing proportion with the remaining genera keeping as ‘other genera’. Each stacked bar plot represents the abundance of bacterial and fungal genera in each sample of the corresponding category. Notable differences in bacterial and fungal populations are those where the taxon is abundant in sweet yogurt samples, and effectively undetected in the sour yogurt samples. The distribution and relative abundance of the bacterial and fungal genera in the study metagenomes are also available in Data S1.

**Fig. 4.**
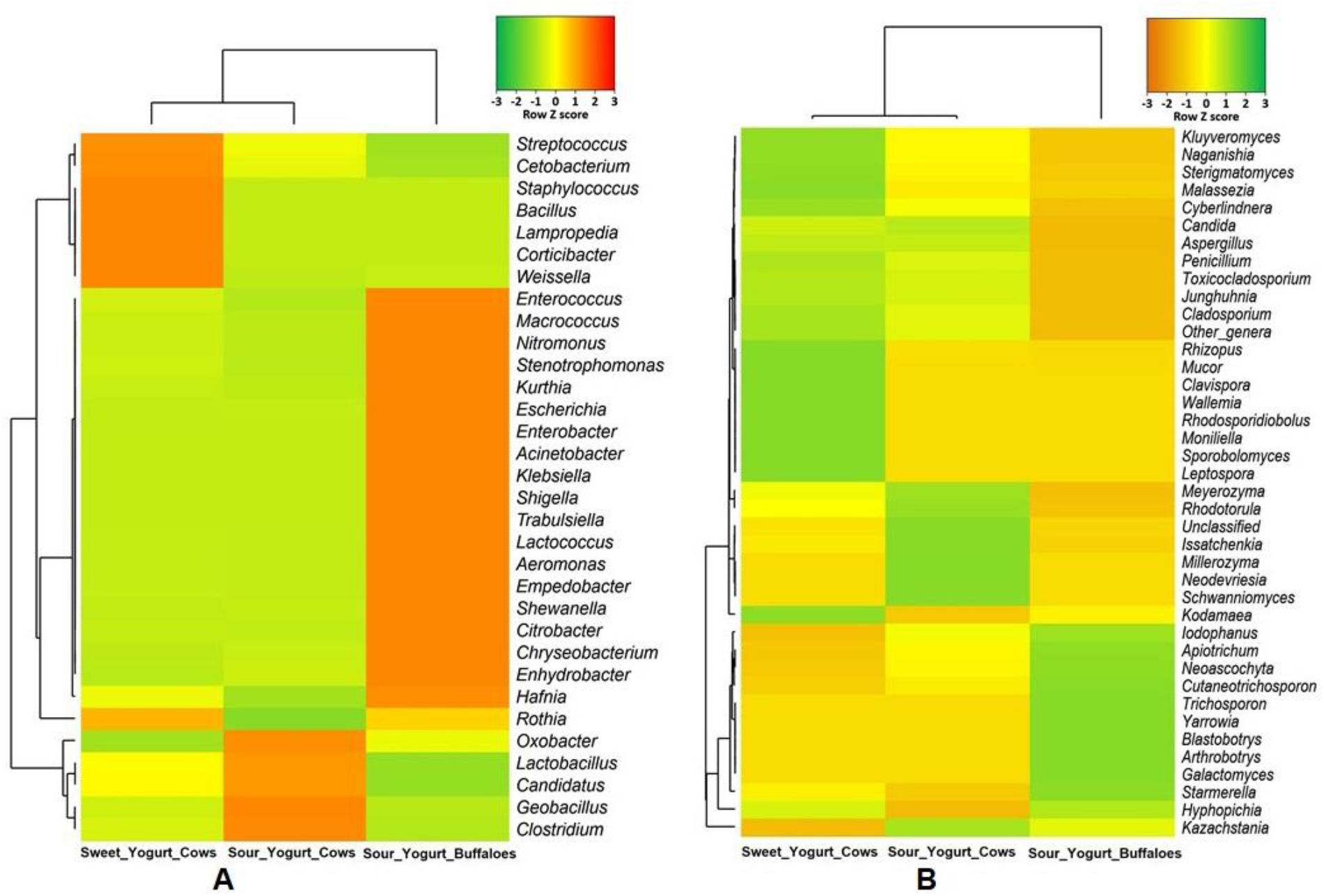
Genus level taxonomic differentiation of bacteria and fungi in sweet yogurt (from cow milk) and sour yogurt of cow and buffalo milk. Heatmap showing 40 top abindant bacterial genera across the sample categories. The color bars (column Z score) at the top represent the relative abundance of each **(A)** bacterial genus and **(B)** fungal genus in the corresponding group. Noteworthy differences in bacterial and fungal populations are those where the genus is abundant in either sweet or sour yogurt samples, and effectively not detected in other metagenomes. The color codes indicate the presence and completeness of each genus in the corresponding sample group, expressed as a value between –3 (lowest abundance) and 3 (highest abundance). The red color indicates the more abundant patterns, while green cells account for less abundant putative genes in that particular metagenome.

**Table 3.**
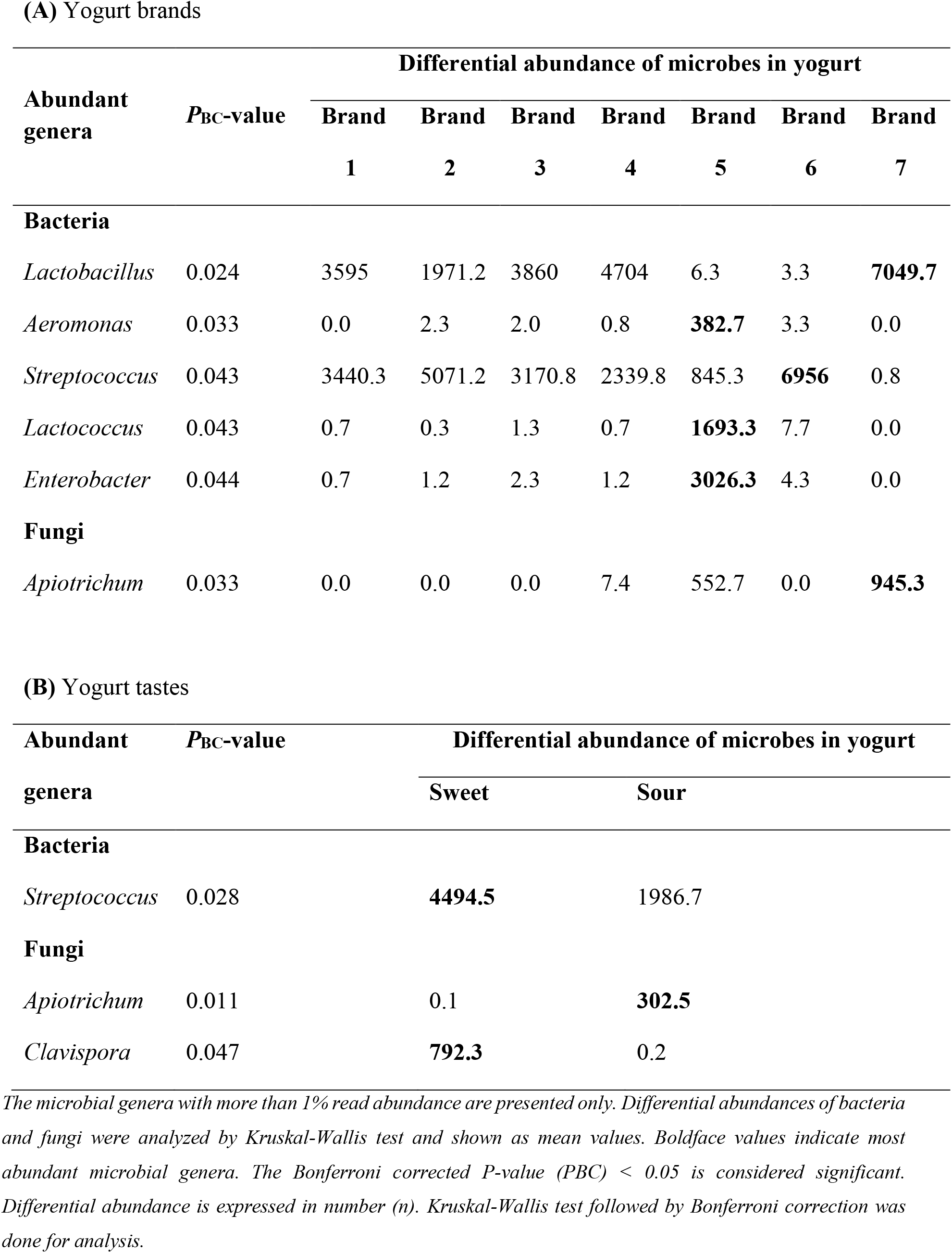
Differential abundance of bacteria and fungi in Bangladeshi yogurt based on brands and tastes.

We further investigated whether genus level relative abundances of the fungi differed between sweet and sour yogurt of different hosts (cow and buffalo) and brands (Fig. 3B, 4B). *Ascomycota* represented more than 98.0% of the total reads at genus level, except for Brand 5 and 6 that consisted of 59.8% of *Basidiomycota* (Fig. S2B). The sweet yogurt metagenome of cow origin had higher relative abundance of *Kluyveromyces* (78.45%), *Clavispora* (12.23%), and *Candida* (5.37%) while *Kluyveromyces* (47.62%), *Trichosporon* (19.74%), *Candida* (8.63%), *Iodophanus* (5.37%), *Apiotrichum* (4.70%), and *Issatchenkia* (2.71%) were the most abundant fungal genera in the sour yogurt metagenome of cow samples (Fig. 3B). Conversely, the sour yogurt sample of buffalo showed higher relative abundances of *Trichosporon* (70.14%), *Iodophanus* (12.07%), *Apiotrichum* (11.94%), and *Neoascochyta* (1.08%), and rest of the fungal genera detected in both sweet and sour yogurt metagenomes had lower relative abundances (<1.0%) (Fig. 3B, Data S1). In addition, similar to bacterial genera, the detected fungal genera showed two distinct clusters in the heatmap representation (Fig. 4B) according to their host origin (i. e. cow and buffalo) rather than tastes of the yogurt. Moreover, the cow originated fungal genera of the yogurt microbiomes were further sub-clustered according to tastes (sweet and sour) (Fig. 4B, Data S1). A differential abundance analysis of the fungal genera showed that higher number of reads (n = 945.3) mapped to *Apiotrichum* in Brand 7 (Table 3A). Similarly, higher abundance of reads assigned for *Clavispora* in sweet brands and *Apiotrichum* in sour brands might reflect the differences in the taste of commercial yogurts. (Table 3B).

To investigate the relationships between relative abundance of microbial genera and presences of minerals in both sweet and sour yogurt, we used non-parametric Kruskal-Wallis tests which revealed significant differences (*P* < 0.05) between the given conditions. Our analysis showed higher level presence of Na, K, Ca and Mg in the sour varieties of SoV3 whereas Cu in the sour yogurt variety of SoV1. Both of these sour varieties (SoV1 and SoV3) were mostly dominated by a single genus of bacteria (*Lactobacillus*; > 97.0%) and fungus (*Kluyveromyces*; 86.23%) (Data S1). By contrast, Fe rich sweet yogurt variety of SwV5 was mainly dominated by *Streptococcus* (89.54%) and *Clavispora* (71.0%). Moreover, the Zn enriched sweet (SwV5) and sour (SoV5) varieties were mostly dominated by both of these bacterial (*Lactobacillus* and *Streptococcus*), and fungal (*Clavispora*) genera were predominantly identified (Data S1).

## Discussion

Yogurt is a popular food in Bangladesh. In spite of the nutritional benefits of the yogurt which is commercially manufactured from bovine and bubaline milk, their consumption and utilization are scarce in many developing countries like Bangladesh. In this context, the production of dairy products such as yogurt using cow and buffalo milk or mixtures of both species of milk could be an interesting and feasible strategy to promote and expand the dairy industry in Bangladesh. Moreover, optimized mixing proportions can also improve the quality of fermented dairy products and develop new ones with specific nutritional (biochemical), physicochemical, sensory and rheological properties. Hence, this research was conducted to explore the nutritional quality assessment, level of mineral constituents or contamination and to explore the hub of microbial consortia in Bangladeshi yogurt of different brands (Brand 1–Brand 7) and taste types (sweet and sour). The biochemical analyses showed that the study samples had a high range of pH (5.28–6.33) which revealed a discordance with the previous report on yogurt pH value (4.6–5.06)^27^. The sour yogurt had comparatively lower pH than the sweet yogurt samples. The control of pH and acidity are undoubtedly important parameters in yogurt processing due to their functional contribution in curd coagulation, ripening, and shelf life. In sour yogurt the decrease of pH could be associated with the fermentation of lactose to lactic acid by lactic acid bacteria (LAB)^28^. Differences in pH values among the samples of two different taste types and seven commercial brands indicates likely influence of the rate of acidification during bacterial fermentation. The pH values of milk during the high heat treatment significantly influenced many properties of the protein particles and their behavior in yogurt fermentation^29^. Moreover, improper incubation time and temperature in addition with regional and source differences might be linked with the high range of pH of the tested yogurt samples.

During this study, the average fat content of yogurt samples ranged from 0.25 to 2.52% (w/w) with no significant differences among the yogurt brands and tastes. Three samples showed individual fat percentages above 3.25, nevertheless, their mean difference didn’t show statistical significance. According to USDA and FDA statements, yogurt is labeled as non-fat, low-fat and regular if it contains less than 0.5%, 0.5–2.0% and at least 3.25% of fat, respectively^30,31^. The nutritive value of yogurt prepared from cow and buffalo milk with respect to MC, TS, SNF and ash ranged between 60–88%, 12–39%, 12–38%, and 0.8–1.25%, respectively. Majority of samples had an average MC below the normal range while having mean TS, SNF and ash that exceeded the average nutritive value^28^. Usually, the sour yogurt samples had higher MC than the sweet samples. Conversely, the mean TS and SNF content remained higher in sweet yogurt samples compared to the sour ones. However, an abnormal increase in the value of TS and SNF (e.g., in SwV4 of Brand 4) substantiates the removal of cream and addition of adulterants. This study did not reveal any significant differences in fat contents of the yogurt samples of both sweet and sour tastes and seven commercial products. Previous studies reported that moisture, TS, SNF, fat, protein and ash contents of the yogurt could vary according to source of milk, manufacturing protocol and storage conditions^28,32,33^. Nonetheless, these variations are in accordance with the differences in approximate composition reported in previous reports^28,33,34^.

Yogurt is gaining more popularity in Bangladesh due to its nutritional properties and potentially beneficial effects in human health. The mineral contents of the yogurt samples of the current study showed distinct variations as evidenced by the higher value for Na content in sweet yogurt samples. Among different minerals, the lowest amount of Na was found in sour yogurt of Brand 4 (SoV43) and Brand 3 (SoV2). The sour yogurt samples possessed a higher mean content of K, Ca and Mg. Variation for the nutritional value of minerals (Na, K, Ca and Mg) and trace elements (Fe and Zn) for low-fat and fat-free yogurt were found in the range 700–770 mg/kg for Na, 2341–2552 mg/kg for K, 1829–1988 mg/kg for Ca, 170–188 mg/kg for Mg, 0.82–0.88 mg/kg for Fe and 5–10 mg/kg for Zn^14^. The imbalance in mineral content might be due to the adulteration of yogurt with unusual compounds^34^. The mineral contents detected in the present study in most of the samples confirmed the role of yogurt as a source of essential nutrients in comparison with raw milk. Moreover, this dairy product could be represented as an excellent alternative to milk for the lactose intolerant population^28,35,36^.

Yogurt is a good source of trace elements such as Fe, Cu and Zn which are essential for human and animal metabolism and growth^37^. We found significant variations in the trace minerals content of the yogurt samples in this study and the mean content of Fe always remained higher in sweet yogurt samples (except for one sour sample; SoV5). The highest accumulation of Zn was recorded in sour yogurt of Brand 7 while the least concentration of Zn was detected in Brand 4 of sweet yogurt. Likewise, the Cu content also varied among different brands of yogurt of two different tastes. The dietary reference intakes according to WHO (World Health Organization) for Zn and Cu are 8–11 and 0.7–0.9 mg/kg, respectively^38^. Only two samples, SwV5 and SoV5 were in the range of permissive value for Zn, while all others were below the range. The accumulation level of Cu in all samples crossed the reference value, while showing conformity among them^28,35,36^.

Microbial diversity of both bacteria and fungi were investigated through amplicon sequencing targeting V3–V4 regions of the 16S rRNA and V9 region of 18S rRNA gene, respectively. In this study, 76 bacterial genera were detected which were mostly represented by increased phylum-level signature of *Firmicutes* (>92.89%) and *Proteobacteria* (>7%). *Streptococcus* and *Lactobacillus* (>50% relative abundances) were detected as the most dominant LAB genera in all brands of sweet and sour varieties of yogurt. Strains of these two bacteria are generally known as the traditional starter bacteria which have been used for manufacturing yogurt since ancient times^39^. Microbial composition and diversity differ from raw to pasteurized milk, and between curd, whey, cheese and yogurt. The raw milk microbiota is influenced by microbes present in the teat canal, the surface of teat skin, hygiene practices, animal handlers, healthy and disease condition of the mammary glands, and the indigenous microbiota of equipment and storage containers^6,9,16^. Remarkably, the sour yogurt samples showed higher taxonomic abundances of *Lactobacillus*, *Streptococcus*, *Macrococcus*, *Enterobacter*, *Empedobacter*, *Lactococcus* and *Aeromonas* genera. This taste specific microbiomes discrimination was more evident in the sour yogurt samples of buffalo since most of the genera had higher relative abundances in buffalo yogurt metagenome. There is growing consensus that the microbiota is an ecosystem working together to keep humans healthy, with no specifically defined structure. The origin of the milk would also seem to influence the levels of diversity therein, with cow milk appearing to be more diverse than that from buffalo, goat and sheep^9,40^. However, until now, no substantial information is available regarding the breed specific association of microbiomes in yogurt samples of different tastes.

We also observed a notable link between *Streptococcus* and sweetness of yogurt (*P*_BC_ = 0.028) in terms of taste of the yogurt showing accordance with recent findings on enzymatic modification of starter bacteria to derive sweetness enhanced yogurt^8^. This might be due to the lower incubation temperature (<40 °C) and shorter incubation period and higher pH along with the addition of sugar that favor the growth of *Streptococcus* over *Lactobacillus*^41^. Pathogenic bacterial genera reported in previous studies including *Pseudomonas*, *Micrococcus*^20^, *Enterococcus*, *Leuconostoc*, *Pediococccus*^21^ were detected at very low levels in this study (Data S1, not shown in Fig.). However, OTU reads assigned for *Aeromonas*, *Enterobacter*, *Lactococcus*, and *Acinetobacter* were found to be higher in Brand 5 while in other brands their relative abundances remained quite insignificant. *Aeromonas* genus exists in soil and aquatic environments, and is responsible for gastrointestinal infection in humans. It is also found to be associated with food-borne diseases like traveler’s diarrhea and water-borne disease outbreaks^42^. Notable that, bacteria comprising *Enterobacter* genus are fecal coliforms associated with bovine intestinal infection that emerge as contaminants in yogurt^43^. Species of the genus *Macrococcus* are naturally widespread commensals primarily isolated from animal skin and dairy products but becoming increasingly recognized as veterinary pathogen^7,44,45^, and thus could have dramatic consequences for public health. Though *Acinetobacter*, gram negative rods are present in the normal flora of human skin, they evolve as opportunistic pathogens and cause disease in immunocompromised patients via food or water^6,46^. *Lactococcus* along with *Acinetobacter* are reported to produce off-flavor, rancidity and ropiness in dairy products during storage^43^. The flavor of the yogurt results from the activity of the microorganisms in the starter cultures. The predominantly identified LAB found to be shared between sweet and sour yogurt samples included different species of *Lactobacillus*, *Streptococcus*, *Enterobacter*, *Lactococcus*, *Aeromonas*, *Acinetobacter*, *Macrococcus* etc. These bacteria are associated with different biochemical conversions like glycolysis, proteolysis and lipolysis to be associated with the production of various flavor and off-flavor compounds for yogurt catalyzed by microbial enzymes^47^.

The present study marks an additional step towards identifying the significant co-occurrence of fungi with bacterial population in yogurt samples. In comparison to bacteria, the relative abundance and diversity of fungi remain slightly lower. Currently, there is no extensive evidence supporting the role of fungi in the fermentation of dairy products like yogurt. We found significant differences in the composition and diversity of the fungal taxa across the sweet and sour yogurt metagenomes. The *Kluyveromyces*, a genus of Ascomycetous yeasts was found as the mostly abundant fungal genus in the metagenomes of both sweet and sour yogurt of cow origin while the sour yogurt of buffalo milk had the highest abundance of *Trichosporon*, a genus of Basidiomycota fungi. Fungal genera predominantly identified in these metagenomes were *Clavispora*, *Candida*, *Iodophanus*, *Apiotrichum*, and *Issatchenkia*. Fungal contamination is a frequent scenario in dairy industries indicating unhygienic practices during manufacturing of these products. Among the previously reported fungal genera *Kodamaea*, *Clavispora*, *Candida*, and *Tricosporon*^20^, the first one was not detected in this research. In addition, OUT reads assigned to detected fungal genera of *Kluyveromyces*, *Trichosporon*, *Candida*, *Clavispora*, and *Apiotrichum* in this study were not reported earlier in Bangladeshi yogurt samples. The ability of the different species of the *Kluyveromyces* genus to metabolize milk constituents (lactose, proteins and fat) makes them very important in cheese and yogurt ripening and fermented milk products as they contribute to maturation and aroma formation^48^. However, despite the importance of fungi in the dairy products, commercial yeast starters like LAB are not in routine use, and fungal flora developing in the yogurt and other dairy products appears as a result of spontaneous contamination.

Yeasts sometimes are involved in lactose fermentation and production of acid, yet they are major spoilage organisms of yogurt, as they are able to grow in low pH, low storage temperature, low water activity, elevated salt concentration, and can even multiply during storage of product. Fungal spoilage generally occurs during final packaging of the finished product with the addition of ingredients and contributes to food-borne diseases in humans. In recent years, several microorganisms including bacteria (e.g., LAB), molds and yeasts are being used in the dairy industry for their functionalities as both starters and secondary flora. This is the first report on nutritional composition and microbial diversity of commercial yogurt of different brands and tastes in Bangladesh.

Minerals are associated with bacterial physiological processes reshaping the gut microbiota’s structure and composition^49^. Fermented dairy products like yogurt are rich in many minerals with a higher bioavailability. The contribution of LAB with particular emphasis to *Streptococcus* and *Lactobacillus* seems to play an important role in the absorption of these minerals (Na, Ca, K, Mg, Cu and Zn), inhibition of other pathogenic microbiota, and in the stimulation of intestinal secretion for digestion of yogurt^50^. Furthermore, fermentation with LAB to produce yogurt results in an acidic environment that can enhance the bioavailability of these minerals^50^. The lower pH of the gut maintains Ca and Mg in their ionic forms, facilitating their enhanced absorption. The pH also facilitates the ligand binding affinity of zinc for transportation across the intestinal wall, resulting in increased absorption of Zn^50^. However, unlike *Saccharomyces*, the role of *Clavispora* with higher mineral contents, and their role in the bioavailability in yogurt needs to be determined *in vivo*. This study will serve as a benchmark for further work on the yogurt microbiome structure and quality manufactured by different companies. Future studies should focus on the contribution of the microbiota in the quality of yogurt and daily yogurt intake to human health in Bangladesh.

## Conclusion

Yogurt, a popular dairy product, is served as a reservoir for good bacteria. It is considered as one of the best sources of probiotics. With a diverse microbial community and collection of minerals, yogurt both sweet and sour comes with health benefits. In this study, the nutritional composition of yogurt and its microbial (bacterial and fungal) communities were investigated by cutting-edge molecular and sequencing technique using 16S and 18S rRNA. We found that the sweet yogurt samples had higher mean pH value, fat, moisture, TS and SNF contents (%, w/w) while the sour samples possessed higher amounts of ash and minerals (Na, K, Ca, Mg Fe, Zn and Cu). The bacterial and fungal diversity were found significant among the brands and between sweet and sour tastes. The metagenomic study showed that *Firmicutes* and *Ascomycota* comprised more than 95% of the read abundances for bacteria and fungi, respectively. The bacterial communities were dominated by *Lactobacillus* and *Streptococcus* in six out of seven brands, except for one sour buffalo yogurt that had significantly higher bacterial diversity and profusion of opportunistic pathogens. *Kluyveromyces*, an Ascomycetous yeast genus, was found as the predominantly abundant fungal genus in six brands of yogurt, however like bacteria, the differences in fungal diversity also found prominent for the sour buffalo yogurt. It is highlighted that higher abundance for *Streptococcus* and *Clavispora* differentiate sweet varieties from sour yogurt which further corroborated with the differences in mineral contents across the sweet and sour varieties of yogurt. The microbiome composition and diversity varied according to the tastes, brands and host-source of the yogurt. This study for the first time demonstrated the nutritional composition and microbiome diversity in yogurt samples of different brands, tastes and hosts which will help the regulatory bodies to monitor yogurt quality and strictly maintain proper guidelines for yogurt manufacturing in Bangladesh.

## Methods

### Sampling

We collected thirty (n = 30) yogurt samples of cow (n = 27) and buffalo (n = 3) milk belonging to seven renowned brands (Brand 1– Brand 7) and two different taste types (sweet; n = 15 and sour; n = 15) from different regions of Bangladesh (Fig. S1, Data S1, Table S1). For this study, seven sampling sites in Bangladesh, i.e. Rangpur, Bogura, Manikgonj, Dhaka, Munshiganj, Chattogram and Cox’s Bazar were selected (Fig. S1) based on the consumer demand of yogurt and its popularity in these selected areas. Aseptic condition was maintained during the collection and transport of products to the laboratory. The samples were randomly collected from the local confectionery shops, placed immediately in a cooling box containing refrigerants (at 4 °C) and transported (within 12 h) to the laboratory. Yogurt samples were stored at –80 °C until further processing for subsequent experiments.

### Nutritional properties of yogurt

The nutritional properties of yogurt samples in terms of biochemical parameters (pH, fat, moisture content, total solid, solid-non-fat, and ash) and mineral contents were determined according to the standard methods. All the biochemical analyses were carried out in triplicate and mean values ± standard deviations (SD) were recorded throughout the article.

#### Determination of physical and biochemical parameters

The pH values of yogurt samples were determined at room temperature using a pH meter (Hanna Instruments, USA). The probe of the pH meter was inserted carefully into the samples and the pH values were measured. The pH measurement was conducted three times for each sample and the average values were recorded. Fat content of yogurt sample was estimated according to the AOAC (Association of Official Analytical Chemists)^29^ by treating the sample with concentrated ammonia solution, then with ethanol, diethyl ether and petroleum ether. The whole process was repeated three times to extract the whole fat content. The solvent was evaporated and the resulting fat sediment was oven-dried (at 100 °C for 2 h) to calculate the percentage (%, w/w) of fat from dry-weight. The moisture content (MC) of a food product is defined as the mass of water present in a known mass of sample. The MC of yogurt was measured following the method described by AOAC^20^. Briefly, the initial weight of the yogurt sample was taken at a constant basis and then oven-dried at 105 °C for 3 h. After over-drying, the sample was immediately placed in a desiccator and the dry (final) weight was again taken. The MC was calculated by subtracting the dry weight from the initial weight of the sample and was expressed in percentage (%, w/w). The amount of total solid (TS) of yogurt was determined by gravimetric method as outlined in AOAC^51^ and was expressed in percentage (%). Solid-non-fat (SNF) is the value of leftovers after fat content is removed from dairy products. It was calculated by subtracting the estimated fat content (%) from TS (%). The ash content was estimated by incinerating the sample in a muffle furnace at 550 °C for 24 h according to the method of AOAC^51^ and was expressed in % (w/w).

#### Determination of mineral contents

After oven-drying and dry-ashing, the inorganic residues left in the crucible were digested with HNO_3_ and diluted up to 100 mL of volume in order to measure the mineral contents of yogurt samples. The diluted samples were then introduced to the Flame Atomic Absorption Spectrophotometer (iCE-3000 FASS, Thermo Fisher Scientific, USA) for the determination of minerals including Na, K, Ca, Mg, Fe, Zn and Cu. The wavelengths used to determine the contents of these minerals were 589.0, 766.5, 422.7, 285.2, 248.3, 213.9, and 324.8 nm, respectively. Standard solutions were prepared at four different concentrations of 0.25, 0.50, 1.0 and 2.0 ppm for all of these elements except Ca for which the concentrations were of 0.5, 1.0, 2.0 and 4.0 ppm. The mineral contents in yogurt samples were estimated spectrophotometrically following the method outlined by Tognato^52^.

### Genomic DNA extraction and amplicon sequencing

The genomic DNA from different yogurt samples was extracted using a commercial microbiome DNA purification kit (Thermo Fisher Scientific, USA) following the manufacturer’s instructions. The amplification of bacterial DNA was achieved by targeting the V3–V4 region of 16S rRNA gene with 30 µL final volume containing 15 µL of 2× master mix (BioLabs, USA), 3 µL of template DNA, 1.5 µL of each V3–V4 forward and reverse primers 341f (5′-CCTACGGGNGGCWGCAG-3′) and 785r (5′-GACTACHVGGGTATCTAATCC-3′), respectively^53^ and the remaining 9 µL of DEPC treated ddH_2_O. A 25 cycle of PCR amplification including initial denaturation at 95 °C for 3 min, denaturation at 95 °C for 30 s, primer annealing at 55 °C for 30 s and elongation at 72°C for 30 s was performed for bacterial DNA with the final extension of 5 min at 72 °C in a thermal cycler (Analytik Jena, Germany).

To amplify fungal DNA, universal primers 1391f (5′-GTAC ACACCGCCCGTC-3′)^54^ and EukBr (5′-TGAT CCTTCTGCAGGTTCACCTAC-3′)^55^ were used targeting V9 hypervariable region of 18S SSU rRNA. PCR mixture for the amplification of fungal DNA was the same as the one used for bacteria. We ran thirty-five cycles of PCR amplification for fungal DNA with the temperature profile of initial denaturation at 94 °C for 3 min, denaturation at 94 °C for 45 s, annealing at 57 °C for 1 min, elongation at 72 °C for 1.5 min and final extension of 10 min at 72 °C. After electrophoresis, the PCR amplicons were visualized in 1.5% agarose gel prepared in 1× TAE buffer. The microbiomes of yogurt were assessed by HTS based on 16S and 18S rRNA genes. We used Agencourt Ampure XP beads (Beckman Coulter, Brea, USA) for PCR products purification, and used the Nextera XT index kit (Illumina, San Diego, USA) for paired-end library preparation according to Illumina standard protocol (Part# 15044223 Rev. B). Paired-end (2×300 bp reads) sequencing of the prepared library pools was performed using MiSeq high throughput kit (v3 kit, 600 cycles) with an Illumina MiSeq platform (Illumina, USA).

### Data processing and downstream bioinformatics

FastQC pipeline was used to examine the primary quality of Illumina sequences of microbiomes^56^. Poor sequencing reads and adapter sequences were trimmed or removed by via BBDuk (with options k = 21, mink = 6, ktrim = r, ftm = 5, qtrim = rl, trimq = 20, minlen = 30, overwrite = true)^44^. MeFiT (merging and filtering tool) was employed to efficiently merge overlapping paired-end sequence reads generated from the Illumina MiSeq with default parameters^57^. Filtering of merged reads, removal of chimeras, *de novo* assembly of reads into operational taxonomic units (OTUs) at 97% similarity level were performed using micca (microbial community analysis) OTU software pipeline (version 1.7.0)^58^. Taxonomic classification of the representative bacterial OTUs was performed using micca classification against SILVA 1.32 release^59^, while fungal OTUs were classified against UNITE database^60^. PASTA algorithm^61^ and FastTree (v2.1.8)^62^ GTR+CAT model were used for multiple sequence alignment (MSA) and phylogenetic tree construction.

To estimate the within sample diversity (α-diversity), we calculated the observed OTUs, Shannon and Simpson diversity indices in microbiomeSeq^63^ and visualized using phyloseq^64^ R package. To visualize differences in microbial diversity across the sample groups, a non-metric multidimensional scaling (NMDS) based on Bray-Curtis dissimilarity of weighted UniFrac metric, and permutational multivariate analysis of variance (PERMANOVA) was performed in Vegan^65^, microbiomeSeq and Phyloseq R packages. Differentially abundant bacteria and fungi at genus level were identified using non-parametric Kruskal-Wallis test followed by Bonferroni correction to exclude 5% false discovery rate (FDR) at 0.05 level of significance in QIIME (v1.9.1)^66^.

### Statistical analysis

All the experiments were carried out in triplicate and the data were expressed in average values ± SD. The statistical analyses were performed using the R package (version i386-4.0.2). Statistical significance in mean differences of various biochemical parameters of yogurt samples was examined using one-way analysis of variance (ANOVA). Post-Hoc test was performed for pair-wise comparison among different brand varieties and within themselves. The results were evaluated statistically with the probability (*P*)-value. At 95% confidence interval (CI), *P* < 0.05 is considered statistically significant. The non-parametric Kruskal-Wallis rank sum test was used to evaluate differences in the relative percent abundance of microbial taxa in different yogurt groups.

## Author contributions

S.M.R.I. and A.M. conceived and designed the study, and monitored laboratory experiments throughout the research; A.Y.T. and M.H.R. collected yogurt samples and performed molecular experiments with the guidance of AMAM. Z.S.; M.J.F. and A.T. executed metagenomic sequencing of amplicons and bioinformatics and statistical analyses of the sequencing data; A.Y.T. and M.A.B. measured the mineral contents of samples; A.Y.T., M.H.R. and S.M.J.H. analyzed other biochemical parameters of yogurt samples; A.Y.T. analyzed the biochemical data, interpreted results, drafted the manuscript and prepared the supplementary information files; M.J.F. and M.N.H. performed the bioinformatics analysis and made visual presentation of data and revised the manuscript draft; S.M.R.I. made critical revision to the manuscript revised; A.Y.T., M.N.H., M.M, A.M., AMAM. Z.S. and A.T. made necessary modifications to the revised version of the manuscript; A.Y.T., A.M. and S.M.R.I. prepared the final version of the manuscript; All authors critically reviewed, read and approved the final manuscript with final check by S.M.R.I.; S.M.R.I. and A.Y.T. contributed equally to this work as first co-authors.

## Data availability

All data analyzed during the present work are included in this article and its supplementary information. The raw sequence data in fastq files are currently available in National Centre for Biotechnology Information (NCBI) under the BioProject accession number PRJNA733702.

## Ethics statement

Not applicable.

## Acknowledgements

This study was partially supported by the University Grants Commission (UGC) of Bangladesh, Dhaka (Research grant-Ref. No: 37.01.0000.073.02.029.19 2717, dated: 22/03/2020). The authors also thank the Department of Genetic Engineering and Biotechnology, Chittagong University; CVASU, BFRI, BCSIR, Chattogram and Helicobacter Research Laboratory, Marshall Centre for Infectious Disease Research and Training, School of Biomedical Sciences, University of Western Australia, Curtin University, Perth, WA, Australia for providing laboratory facilities to make this study possible.

## Supplementary information

Supplementary information supporting the results of the study are available in this article as Figs. S1, S2, Tables S1, S2 and Data S1.

## Reference

1 Astrup, A. Yogurt and dairy product consumption to prevent cardiometabolic diseases: epidemiologic and experimental studies. The American journal of clinical nutrition 99, 1235S–1242S (2014).

2 González, S. et al. Fermented dairy foods: impact on intestinal microbiota and health-linked biomarkers. Frontiers in microbiology 10, 1046 (2019).

3 Sivamaruthi, B. S., Kesika, P. & Chaiyasut, C. Impact of fermented foods on human cognitive function—A review of outcome of clinical trials. Scientia pharmaceutica 86, 22 (2018).

4 Chandan, R. C. & Kilara, A. Manufacturing yogurt and fermented milks. (Wiley Online Library, 2013).

5 Warinner, C. et al. Direct evidence of milk consumption from ancient human dental calculus. Scientific reports 4, 1–6 (2014).

6 Hoque, M. N. et al. Metagenomic deep sequencing reveals association of microbiome signature with functional biases in bovine mastitis. Scientific reports 9, 1–14 (2019).

7 Hoque, M. N. et al. Microbiome dynamics and genomic determinants of bovine mastitis. Genomics 112, 5188–5203 (2020).

8 Luzzi, G., Brinks, E., Fritsche, J. & Franz, C. M. Microbial composition of sweetness-enhanced yoghurt during fermentation and storage. AMB Express 10, 1–7 (2020).

9 Yeluri Jonnala, B., McSweeney, P. L., Sheehan, J. J. & Cotter, P. D. Sequencing of the cheese microbiome and its relevance to industry. Frontiers in microbiology 9, 1020 (2018).

10 Pei, R., Martin, D. A., DiMarco, D. M. & Bolling, B. W. Evidence for the effects of yogurt on gut health and obesity. Critical reviews in food science and nutrition 57, 1569–1583 (2017).

11 Yilmaz-Ersan, L. & Kurdal, E. The production of set-type-bio-yoghurt with commercial probiotic culture. International Journal of Chemical Engineering and Applications 5, 402 (2014).

12 Muehlhoff, E., Bennett, A. & McMahon, D. Milk and dairy products in human nutrition. (Food and Agriculture Organization of the United Nations (FAO), 2013).

13 Park, Y. W., Haenlein, G. & Ag, D. S. Milk and dairy products in human nutrition. Wilet-Blackwell. A John Wiley & Sons, Ltd., Publication 700 (2013).

14 Raw, P. Composition of Foods Raw, Processed, Prepared USDA National Nutrient Database for Standard Reference, Release 25. United States Department of Agriculture (USDA) (2012).

15 Chandan, R. C. Dairy processing and quality assurance: an overview. Dairy Process Quality Assur, 1–40 (2008).

16 Hoque, M., Sultana, M. & Hossain, A. Dynamic Changes in Microbiome Composition and Genomic Functional Potentials in Bovine Mastitis. J Data Mining Genomics Proteomics 12, 232 (2021).

17 Walsh, A. M. et al. Strain-level metagenomic analysis of the fermented dairy beverage nunu highlights potential food safety risks. Applied and environmental microbiology 83 (2017).

18 Sulaiman, I. M., Jacobs, E., Simpson, S. & Kerdahi, K. Molecular identification of isolated fungi from unopened containers of Greek yogurt by DNA sequencing of internal transcribed spacer region. Pathogens 3, 499–509 (2014).

19 Shangpliang, H. N. J., Rai, R., Keisam, S., Jeyaram, K. & Tamang, J. P. Bacterial community in naturally fermented milk products of Arunachal Pradesh and Sikkim of India analysed by high-throughput amplicon sequencing. Scientific reports 8, 1–10 (2018).

20 Nahidul-Islam, S. M., Kuda, T., Takahashi, H. & Kimura, B. Bacterial and fungal microbiota in traditional Bangladeshi fermented milk products analysed by culture-dependent and culture-independent methods. Food Research International 111, 431–437 (2018).

21 Harun-ur-Rashid, M., Togo, K., Ueda, M. & Miyamoto, T. Identification and characterization of dominant lactic acid bacteria isolated from traditional fermented milk Dahi in Bangladesh. World Journal of Microbiology and Biotechnology 23, 125–133 (2007).

22 Meshref, A. M., Moselhy, W. A. & Hassan, N. E.-H. Y. Heavy metals and trace elements levels in milk and milk products. Journal of food measurement and characterization 8, 381–388 (2014).

23 Pšenková, M., Toman, R. & Tančin, V. Concentrations of toxic metals and essential elements in raw cow milk from areas with potentially undisturbed and highly disturbed environment in Slovakia. Environmental Science and Pollution Research 27, 26763–26772 (2020).

24 Coni, E., Bocca, A., Ianni, D. & Caroli, S. Preliminary evaluation of the factors influencing the trace element content of milk and dairy products. Food chemistry 52, 123–130 (1995).

25 Gaucheron, F. Milk minerals, trace elements, and macroelements. Milk and dairy products in human nutrition: Production, composition and health, 172–199 (2013).

26 Elbagermi, M., Alajtal, A. & Edwards, H. A comparative study on the physicochemical parameters and trace elements in raw milk samples collected from Misurata-Libya. SOP transactions on analytical chemistry 1, 15–23 (2014).

27 Ali, M., Islam, M., Alam, M. & Islam, M. Quality of yogurt (Dahi) made in laboratory and available in the market of Mymensingh Town in Bangladesh. Pak. J. Biol. Sci 5, 343–345 (2002).

28 Boukria, O. et al. Biochemical, Physicochemical and Sensory Properties of Yoghurts Made from Mixing Milks of Different Mammalian Species. Foods 9 (2020).

29 Ozcan, T., Horne, D. S. & Lucey, J. A. Yogurt made from milk heated at different pH values. Journal of Dairy Science 98, 6749–6758 (2015).

30 Weerathilake, W., Rasika, D., Ruwanmali, J. & Munasinghe, M. The evolution, processing, varieties and health benefits of yogurt. International Journal of Scientific and Research Publications 4, 1–10 (2014).

31 Oludara, O. M. & Bamidele, O. O. in 2019 ASABE Annual International Meeting. 1 (American Society of Agricultural and Biological Engineers).

32 Clark, S. & García, M. B. M. A 100-year review: Advances in goat milk research. Journal of Dairy Science 100, 10026–10044 (2017).

33 Aswal, P., Shukla, A. & Priyadarshi, S. Yoghurt: Preparation, characteristics and recent advancements. Cibtech Journal of Bio-Protocols 1, 32–44 (2012).

34 Karami, M. Investigation of physicochemical, microbiological, and rheological properties and volatile compounds of ewe and cow milk yoghurt. Journal of Agricultural Science and Technology 20, 1149–1160 (2018).

35 de la Fuente, M. A., Montes, F., Guerrero, G. & Juárez, M. Total and soluble contents of calcium, magnesium, phosphorus and zinc in yoghurts. Food Chemistry 80, 573–578 (2003).

36 Güler, Z. Levels of 24 minerals in local goat milk, its strained yoghurt and salted yoghurt (tuzlu yoğurt). Small Ruminant Research 71, 130–137 (2007).

37 Stawarz, R., Formicki, G. & Massanyi, P. Daily fluctuations and distribution of xenobiotics, nutritional and biogenic elements in human milk in Southern Poland. Journal of Environmental Science and Health, Part A 42, 1169–1175 (2007).

38 Organization, W. H. Trace elements in human nutrition and health. (World Health Organization, 1996).

39 Tamime, A. & Deeth, H. Yogurt: technology and biochemistry. Journal of food protection 43, 939–977 (1980).

40 Quigley, L. et al. High-throughput sequencing for detection of subpopulations of bacteria not previously associated with artisanal cheeses. Applied and environmental microbiology 78, 5717–5723 (2012).

41 Chandan, R. C., Kilara, A. & Shah, N. P. Dairy processing and quality assurance. (John Wiley & Sons, 2009).

42 Igbinosa, I. H., Igumbor, E. U., Aghdasi, F., Tom, M. & Okoh, A. I. Emerging Aeromonas species infections and their significance in public health. The Scientific World Journal 2012 (2012).

43 Lu, M. & Wang, N. S. in The microbiological quality of food 151–178 (Elsevier, 2017).

44 Hoque, M. N. et al. Insights into the resistome of bovine clinical mastitis microbiome, a key factor in disease complication. Frontiers in Microbiology 11, 860 (2020).

45 MacFadyen, A. C., Fisher, E. A., Costa, B., Cullen, C. & Paterson, G. K. Genome analysis of methicillin resistance in Macrococcus caseolyticus from dairy cattle in England and Wales. Microbial genomics 4 (2018).

46 de Amorim, A. M. B. & dos Santos Nascimento, J. Acinetobacter: an underrated foodborne pathogen? The Journal of Infection in Developing Countries 11, 111–114 (2017).

47 Chen, C. et al. Role of lactic acid bacteria on the yogurt flavour: A review. International Journal of Food Properties 20, S316–S330 (2017).

48 Awasti, N. & Anand, S. in Dairy Processing: Advanced Research to Applications 243–262 (Springer, 2020).

49 García-Legorreta, A. et al. Effect of dietary magnesium content on intestinal microbiota of rats. Nutrients 12, 2889 (2020).

50 El-Abbadi, N. H., Dao, M. C. & Meydani, S. N. Yogurt: role in healthy and active aging. The American journal of clinical nutrition 99, 1263S–1270S (2014).

51 Horwitz, W., Chichilo, P. & Reynolds, H. Official methods of analysis of the Association of Official Analytical Chemists. Official methods of analysis of the Association of Official Analytical Chemists. (1970).

52 Tognato, E. Characterization of major mineral contents in milk of four cattle breeds. (2015).

53 Klindworth, A. et al. Evaluation of general 16S ribosomal RNA gene PCR primers for classical and next-generation sequencing-based diversity studies. Nucleic acids research 41, e1–e1 (2013).

54 Ludwig, W. Nucleic acid techniques in bacterial systematics and identification. International journal of food microbiology 120, 225–236 (2007).

55 Devereux, R. & Willis, S. G. in Molecular microbial ecology manual 277–287 (Springer, 1995).

56 Li, X., Nair, A., Wang, S. & Wang, L. in RNA Bioinformatics 137–146 (Springer, 2015).

57 Parikh, H. I., Koparde, V. N., Bradley, S. P., Buck, G. A. & Sheth, N. U. MeFiT: merging and filtering tool for illumina paired-end reads for 16S rRNA amplicon sequencing. BMC bioinformatics 17, 1–6 (2016).

58 Albanese, D., Fontana, P., De Filippo, C., Cavalieri, D. & Donati, C. MICCA: a complete and accurate software for taxonomic profiling of metagenomic data. Scientific reports 5, 1–7 (2015).

59 Quast, C. et al. The SILVA ribosomal RNA gene database project: improved data processing and web-based tools. Nucleic acids research 41, D590–D596 (2012).

60 Nilsson, R. H. et al. The UNITE database for molecular identification of fungi: handling dark taxa and parallel taxonomic classifications. Nucleic acids research 47, D259–D264 (2019).

61 Mirarab, S. et al. PASTA: ultra-large multiple sequence alignment for nucleotide and amino-acid sequences. Journal of Computational Biology 22, 377–386 (2015).

62 Price, M. N., Dehal, P. S. & Arkin, A. P. FastTree 2–approximately maximum-likelihood trees for large alignments. PloS one 5, e9490 (2010).

63 Ssekagiri, A., Sloan, W. & Ijaz, U. (2018).

64 McMurdie, P. J. & Holmes, S. phyloseq: an R package for reproducible interactive analysis and graphics of microbiome census data. PloS one 8, e61217 (2013).

65 Dixon, P. VEGAN, a package of R functions for community ecology. Journal of Vegetation Science 14, 927–930 (2003).

66 Caporaso, J. G. et al. QIIME allows analysis of high-throughput community sequencing data. Nature methods 7, 335–336 (2010).

