## Supplementary material for "Insights into the nutritional properties and microbiome diversity in sweet and sour yogurt manufactured in Bangladesh": Table S1 (from another version of this preprint: version 1, 2021-08-15)

**Table S1.** PERMANOVA of beta-dispersion for bacterial and fungal diversity in Bangladeshi yogurt of different brands and tastes.

| **Bacteria** | |  | **Fungi** | |
| --- | --- | --- | --- | --- |
| **Yogurt brands** | ***P*_permanova_ -value** |  | **Yogurt brands** | ***P*_permanova_ -value** |
| Brand 2, Brand 5 | 6e-06*** |  | Brand 3, Brand 6 | 1e-05*** |
| Brand 2, Brand 7 | 1e-05*** |  | Brand 3, Brand 5 | 3e-05*** |
| Brand 2, Brand 6 | 8e-05*** |  | Brand 2, Brand 3 | 3e-05*** |
| Brand 3, Brand 5 | 1e-03*** |  | Brand 3, Brand 7 | 1e-04*** |
| Brand 3, Brand 7 | 2e-03** |  | Brand 2, Brand 4 | 6e-03** |
| Brand 2, Brand 4 | 2e-03** |  | Brand 4, Brand 6 | 7e-03** |
| Brand 4, Brand 5 | 2e-03** |  | Brand 4, Brand 5 | 1e-02* |
| Brand 3, Brand 6 | 4e-03** |  | Brand 4, Brand 7 | 3e-02* |
| Brand 4, Brand 7 | 5e-03** |  | Brand 1, Brand 2 | 3e-02* |
| Brand 4, Brand 6 | 2e-02* |  | Brand 1, Brand 6 | 3e-02* |
| **Yogurt tastes** | ***P*_permanova_ -value** |  | **Yogurt tastes** | ***P*_permanova_ -value** |
| Sweet – Sour | 0.016 |  | Sweet – Sour | 0.013 |

*Significance level of Ppermanova -values are depicted as P < 0.01 = *, P < 0.001 = ** and P < 0.0001 = ***.*
