## Supplementary material for "Insights into the nutritional properties and microbiome diversity in sweet and sour yogurt manufactured in Bangladesh": Table S2 (from another version of this preprint: version 1, 2021-08-15)

**Table S2.** Taxonomic distribution of bacteria and fungi in yogurt of different tastes, source variety and brands.

| **Taxonomic rank** |  | **Taste and source variety of yogurt** | | |
| --- | --- | --- | --- | --- |
|  |  | **Sweet yogurt cow** | **Sour yogurt cow** | **Sour yogurt buffalo** |
| **Bacteria** |  |  |  |  |
| Phyla (n=11) |  | 10 | 10 | 9 |
| Order (n=30) |  | 24 | 25 | 21 |
| Genus (n=76) |  | 57 | 61 | 51 |
| **Fungi** |  |  |  |  |
| Phyla (n=5) |  | 5 | 4 | 3 |
| Order (n=28) |  | 23 | 24 | 14 |
| Genus (n=70) |  | 43 | 54 | 25 |
